# A reverse chemical ecology approach to explore wood natural durability

**DOI:** 10.1101/779629

**Authors:** Perrot Thomas, Salzet Guillaume, Amusant Nadine, Beauchene Jacques, Gérardin Philippe, Dumarçay Stéphane, Sormani Rodnay, Morel-Rouhier Mélanie, Gelhaye Eric

## Abstract

The natural durability of wood species, defined as their inherent resistance to wood-destroying agents is a complex phenomenon depending of many biotic and abiotic factors. Besides the presence of recalcitrant polymers, the presence of compounds with antimicrobial properties is known to be important to explain wood durability. Based on the advancement in our understanding of fungal detoxification systems, a reverse chemical ecology approach was proposed to explore wood natural durability. A set of six glutathione transferases from the white-rot *Trametes versicolor* was used as targets to test wood extracts from seventeen French Guiana neotropical species. Fluorescent thermal shift assays allowed to quantify interactions between fungal glutathione transferases and these extracts. From these data, a model combining this approach and wood density predicts significantly wood natural durability of the tested species previously estimated by long-term soil bed tests. Overall, our findings confirm that detoxification systems could be used to explore the chemical environment encountered by wood decaying fungi and then wood natural durability.

## Introduction

Wood biodegradation, an essential step in carbon recycling, is a complex phenomenon which depends of many abiotic and biotic factors occurring at different spatial and temporal scales. Wood natural durability, defined as the natural resistance of wood against biologic degradation (Taylor *et al*., 2002) varies according to wood species, geographic regions, and to variations of environmental exposure conditions during trees life. Nevertheless, main wood intrinsic physicochemical properties are essential to wood decay resistance. For instance, wood density, which is widely used as a functional trait in the field of functional ecology has been correlated to wood natural durability (Lehnenbach *et al*., 2019, Beauchène, 2012). Wood components such as extractives are also known to be involved in the wood resistance against decay (Valette *et al.*, 2017). These molecules are not covalently linked to cell walls and could be then extracted using several solvents. A part of these wood extracts possesses antimicrobial and insecticidal activities explaining their involvement in the wood durability (Amusant *et al*., 2014; Rodrigues *et al*., 2011).

In the other side, in forest ecosystems, wood degradation is mainly mediated by specialised microbial communities and in particular by wood decaying fungi. These fungi have evolved to efficiently breakdown and mineralize wood components. In the last few years, comparative genomic studies, confirming previous biochemical and microbiological approaches, have demonstrated the presence of specialized extracellular and intracellular enzymatic networks in these organisms (Nagy *et al.*, 2017). Extracellular networks gather oxidative and hydrolytic enzymes, which catalyse synergistically efficient breakdown of wood polymers. In particular white-rot fungi possess fungal class II peroxidases, which are involved in the lignin breakdown (Floudas *et al.*, 2012). Intracellular networks are mainly involved in the import and catabolism of the degraded wood products and in the detoxification of toxic molecules initially present or generated during wood degradation (Morel *et al.*, 2013; Nagy *et al.*, 2017). Among their extended detoxification system, wood decaying fungi usually possess a larger set of genes encoding glutathione transferases (GSTs). These enzymes are involved in the second step (conjugation) of the detoxification pathways (Morel *et al.*, 2009; Morel et al., 2013) and largely used as indicators of the stress responses in various organisms (Bouzahouane *et al.*, 2018; Bass and Field, 2011, Fernandez Gonzalez *et al.*, 2018). Moreover, GSTs exhibit ligandin properties allowing their non-catalytic interactions with potentially toxic wood molecules (Mathieu *et al.*, 2012).

Molecular mechanisms governing wood durability remain largely to be unravelled and could be explored through the adaptation of organisms involved in wood decay. Since last few years, it was suggested that fungal detoxification systems could give insights about the chemical environment encountered by wood decaying fungi (Deroy *et al.*, 2015; Perrot *et al.*, 2018). In this paper, we propose a reverse chemical approach-like to explore this hypothesis. Chemical ecology is defined by Leal “as the study of the chemical languages, cues, and mechanisms controlling interactions among living beings, including communication among individuals of the same species and between organisms and their environment” (Leal, 2017). From the molecular knowledge of olfaction systems, reverse chemical ecology approaches have been developed to study the behavioural active compounds of various organisms and in particular of insects (Zhu *et al.*,2017; Choo *et al.*, 2018). For instance, interactions between odorant-binding proteins and ligands have been used to screen potential semiochemicals (Li *et al.*, 2018). In the context of wood durability, detoxification systems could be used in a similar approach. Previous studies have indeed demonstrated that GSTs of wood decaying fungi could be used as molecular targets to identify wood molecules with antioxidative and antimicrobial properties (Schwartz *et al.*, 2018, Perrot *et al.*, 2018).

To test the hypothesis that GSTs could be used as indicators of wood durability, a reverse chemical ecology approach was developed using a set of enzymes from the world widespread white-rot fungus *Trametes versicolor* and wood extracts from neotropical forest of French Guiana. The obtained results support the initial hypothesis and demonstrate that such reverse chemical ecology approach could be useful to predict wood durability.

## Results and discussion

### Interactions between GSTs and wood extracts

To set up the experimental design, heartwoods of 17 species from French Guiana tropical forest have been selected (Table 1). Data obtained the heartwoods from *Andira coriacea, Bagassa guianensis, Dicorynia guianensis, Hymenaea courbaril, Peltogyne venosa, Sextonia rubra, and Tabebuia serratifolia* have been previously published (Perrot et al., 2018). Heartwoods from *Abarema jupunba, Bocoa prouacensis, Hirtella bicornis, Oxandra asbeckii, Parkia nitida, Parkia pendula, Pouteria decorticans, Protium gallicum, Swartzia canescens, Vouacapoua americana* were from commercial origin (Degrad Saramaca’s sawmill, Kourou, French Guiana) or harvested as described in Lehnenbach *et al*., 2019. All these woods belong to the DEGRAD database (Beauchène, 2012). The DEGRAD database contains wood density and wood durability data for more than 300 tree species from French Guiana tree species Among the 17 chosen woods, four are classified as very durable (x<10%, x being the relative mass loss obtained during the soil tests), five as durable (10%<x<25%), five moderately durable (25%<x<45%) and three non-durable (x>45%) (Table 1). From the 17 selected species, molecules from corresponding heartwoods have been sequentially extracted using four solvents exhibiting different polarities (dichloromethane, acetone, toluene/ethanol, and water). From this step we obtained a collection of 68 wood extracts.

**Table 1:**
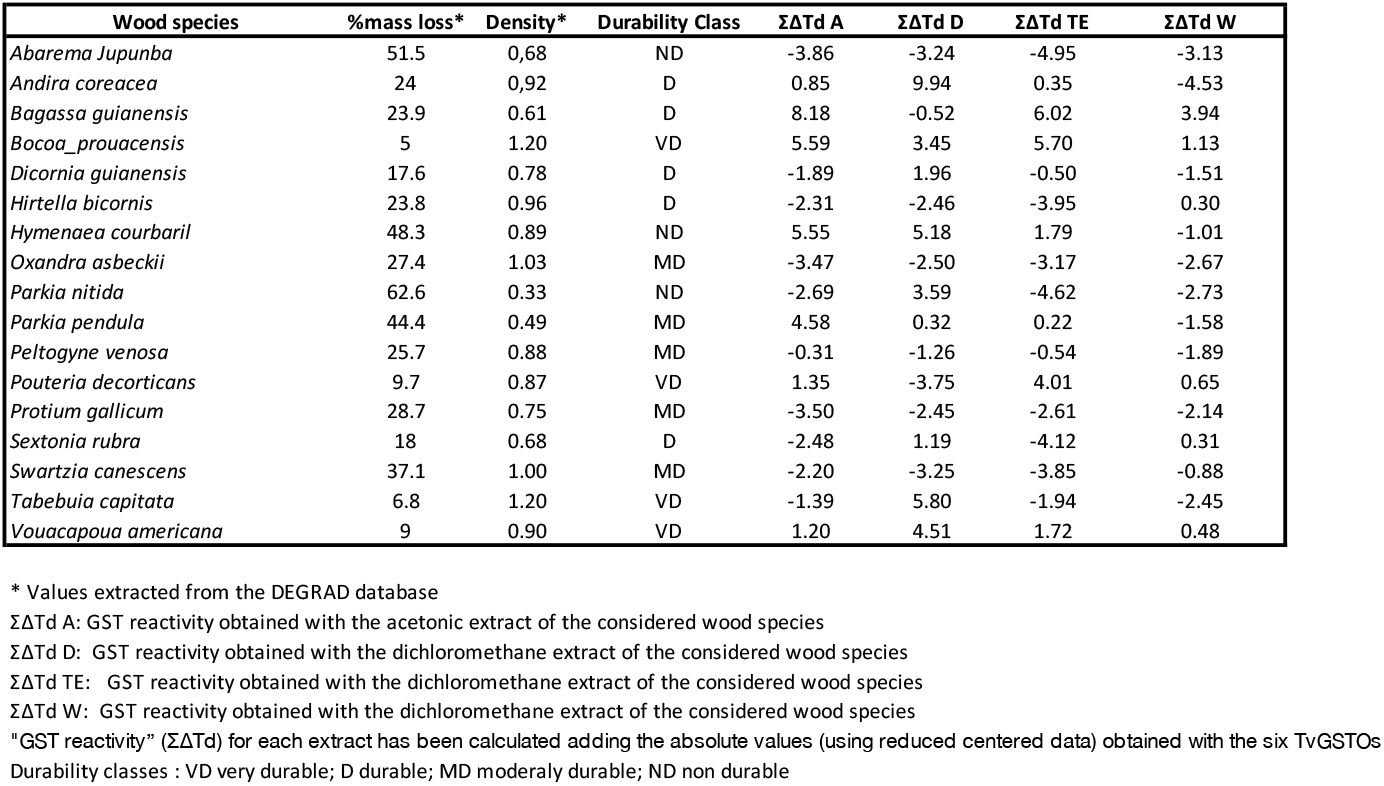
Wood durability (%mass loss), wood density and GST reactivity obtained from the 17 woods of French Guiana Forest.

On the other hand, 6 GSTs belonging to the omega class from *Trametes versicolor* (TvGSTOs), have been used to characterize the wood extracts. Previously, those TvGSTOs have been shown to be able to bind wood polyphenolic compounds (Schwartz *et al.*, 2018; Perrot *et al.*, 2018). The interactions between these TvGSTOs and their ligands were quantitatively measured through a thermal shift assay. This assay allows to determine modification of the protein thermal stability (ΔTd) due to the ligand binding (Deroy *et al*., 2015; Schwartz *et al*., 2018).

Using this approach, interactions between the 68 extracts and the 6 TvGSTOs were then followed (Sup Table 1). Each TvGSTO exhibits a specific pattern of interactions with the tested extracts. For further analysis, a “GST reactivity” value (ΣΔTd) has been calculated for each extract, adding the ΔTd absolute values (using reduced centred data) obtained with the six TvGSTOs (Table 1, supTable 1). Each tested wood was then defined by four values corresponding to the sum of interactions between TvGSTOs and the mixtures obtained after extraction with dichloromethane (ΣΔTdD), acetone (ΣΔTdA); Toluene/Ethanol (ΣΔTdTE) and water (ΣΔTdW). Significant correlations (p<0.05) between three variables ΣΔTdA, ΣΔTdTE and ΣΔTdW were found (Table 2).

**Table 2 :** Correlation (Pearson coefficient) between GST reactivities, wood durability (%mass loss) and wood density.

| Variables | %mass loss | Wood density | ΣΔTd A | ΣΔTdD | ΣΔTd TE | ΣΔTd W |
| --- | --- | --- | --- | --- | --- | --- |
| **%mass loss** | **1** | **-0,657**<br>(p <0.004) | -0,144 | -0,129 | -0,464 | -0,398 |
| **Wood density** |  | **1** | 0,009 | 0,142 | 0,205 | 0,012 |
| **ΣΔTd A** |  |  | **1** | 0,296 | **0,881**<br>(p < 0.0001) | **0,585**<br>(p<0.014) |
| **ΣΔTdD** |  |  |  | **1** | 0,232 | -0,252 |
| **ΣΔTd TE** |  |  |  |  | **1** | **0,604**<br>(p<0.010) |
| **ΣΔTd W** |  |  |  |  |  | **1** |

Values in bold are significant (p<0.05)

### GSTs and wood durability

Wood durability is usually estimated from long term soil bed tests (Meyer *et al*., 2014; XP CEN/TS 15083-2. 2006.). To constitute the DEGRAD database, soil tests have been performed incubating wood blocks in French Guiana soils in 2010/2012 in controlled laboratory conditions as mentioned in Amusant et al. (2014). Mass losses (%) have been measured after 6 incubation months. Data from DEGRAD database were used to quantify wood durability of the seventeen species used in this experiment. 68% of the variability (p<0.006) of the measured mass losses could be explained by a model (linear regression) set up from the four “GST reactivity variables” as shown in Table 2 (%mass loss = 19 + 6.6 ΣΔTdA − 1.6 ΣΔTdD − 6.3 ΣΔTdTE − 4.5 ΣΔTdW; R^2^ = 0.679, p < 0.006). This significant correlation supported the hypothesis that “GST reactivity” could reflect at least partially the wood durability of the considered species.

### GST reactivity and wood density

Wood durability is known to be inversely correlated to wood density (WD). Using more than 300 species of the DEGRAD database, significant correlation between wood density (12% of humidity) and durability (mass losses (%) obtained from soil tests) has indeed been previously observed (R^2^ = 0.43) (Chave *et al*., 2009; Larjavaara *et al*. 2010). We postulated that “GST reactivity” and WD should be link to explain the durability of the tested species. From the same data set (17 heartwoods; Table 1), WD and “GST reactivity” are not significantly correlated (p>0.05). In contrast, as expected, WD could be an indicator of wood durability estimated by soil bed tests (r=−0.657, p= 0.004). Using a linear regression, a model was then constructed using both “GST reactivity” and WD. This model explained more than 81% (p<0.006) of the mass loss variability [% predicted mass loss = 45 – (30 * WD) + (5*ΣΔTdA) − (1,3*ΣΔTdD) − (4,6*ΣΔTdTE) − (4,5*ΣΔTdW)] (Figure 1) demonstrating that “GST reactivity” and WD predict together efficiently wood durability in the used data set. This 2 components model of wood durability reflects woody polymer organisation and the chemical toxicity of the wood.

**Figure 1:**
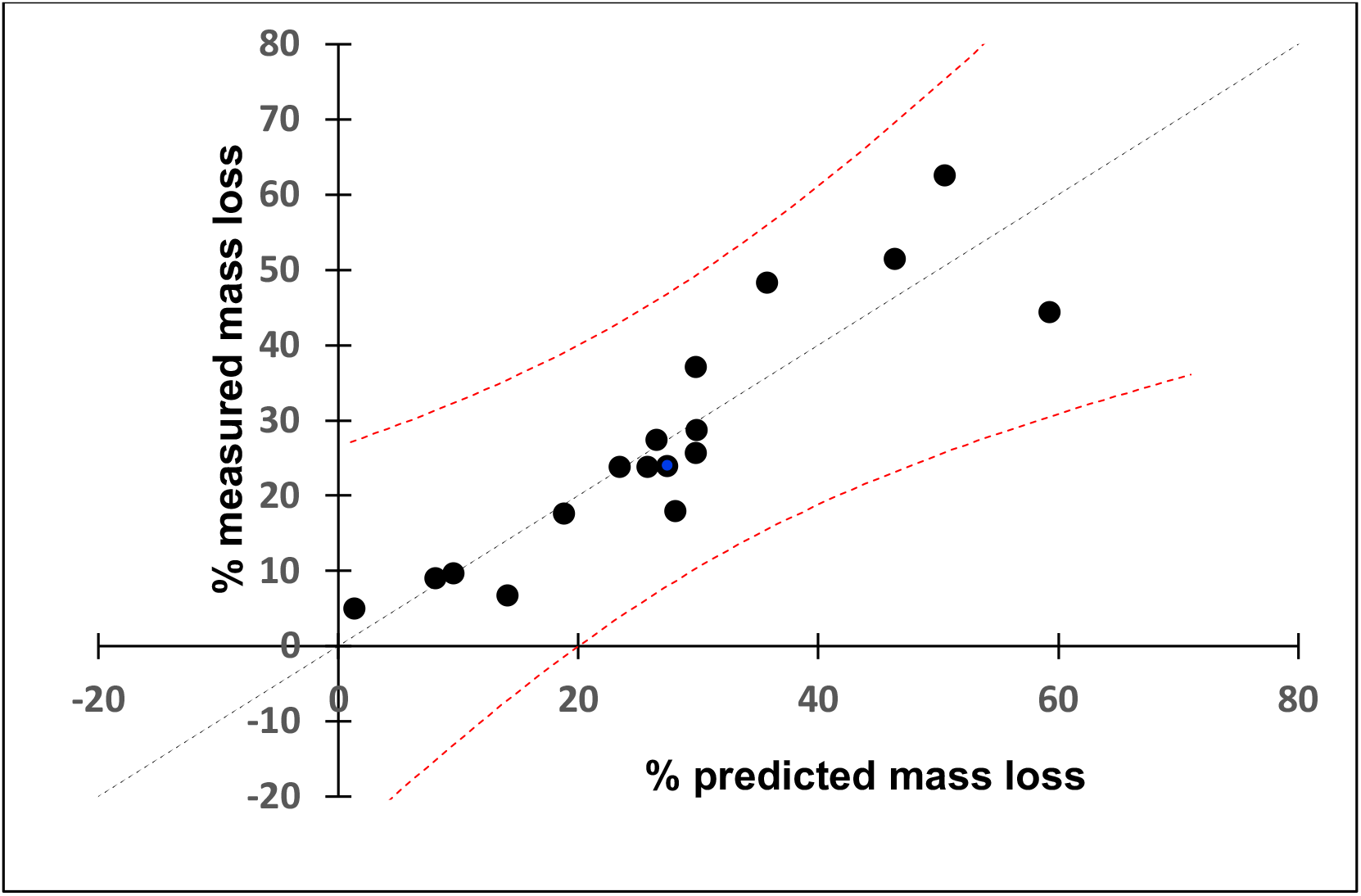
Wood durability model set-up from GST reactivities and wood density (WD). The multiple linear regression model was set up using Xlstat given the following equation : % predicted mass loss = 45 – [30 * WD] + [5*ΣΔTdA - 1,3*ΣΔTdD - 4,6*ΣΔTdTE - 4,5*ΣΔTdW]; R^2^ = 0.818, p<0.006.

Besides macro and micro-environmental factors, wood intrinsic properties trigger its microbial degradation (Amusant *et al*., 2014; Valette *et al*., 2018). Wood natural durability is due to the presence and organization of the recalcitrant polymers (cellulose, hemicellulose and lignin) and also of antimicrobial molecules (wood extracts) (Valette *et al*., 2017). Based on the molecular characterization of the detoxification systems found in wood decaying fungi, we propose here a new reverse chemical ecology approach to study this complex phenomenon. Combining such approach and wood density measurement should give new perspectives for studying wood degradation in various ecosystems.

## Acknowledgements

This work was supported by a grant overseen by the French National Research Agency (ANR) as part of the “Investissements d’Avenir”program (ANR-11-LABX-0002-01, Lab of Excellence ARBRE) and the Region Lorraine Research Council. We thank Solène Telliez and Fanny Saiag for technical assistance.

---

**Supplemental Table 1.**
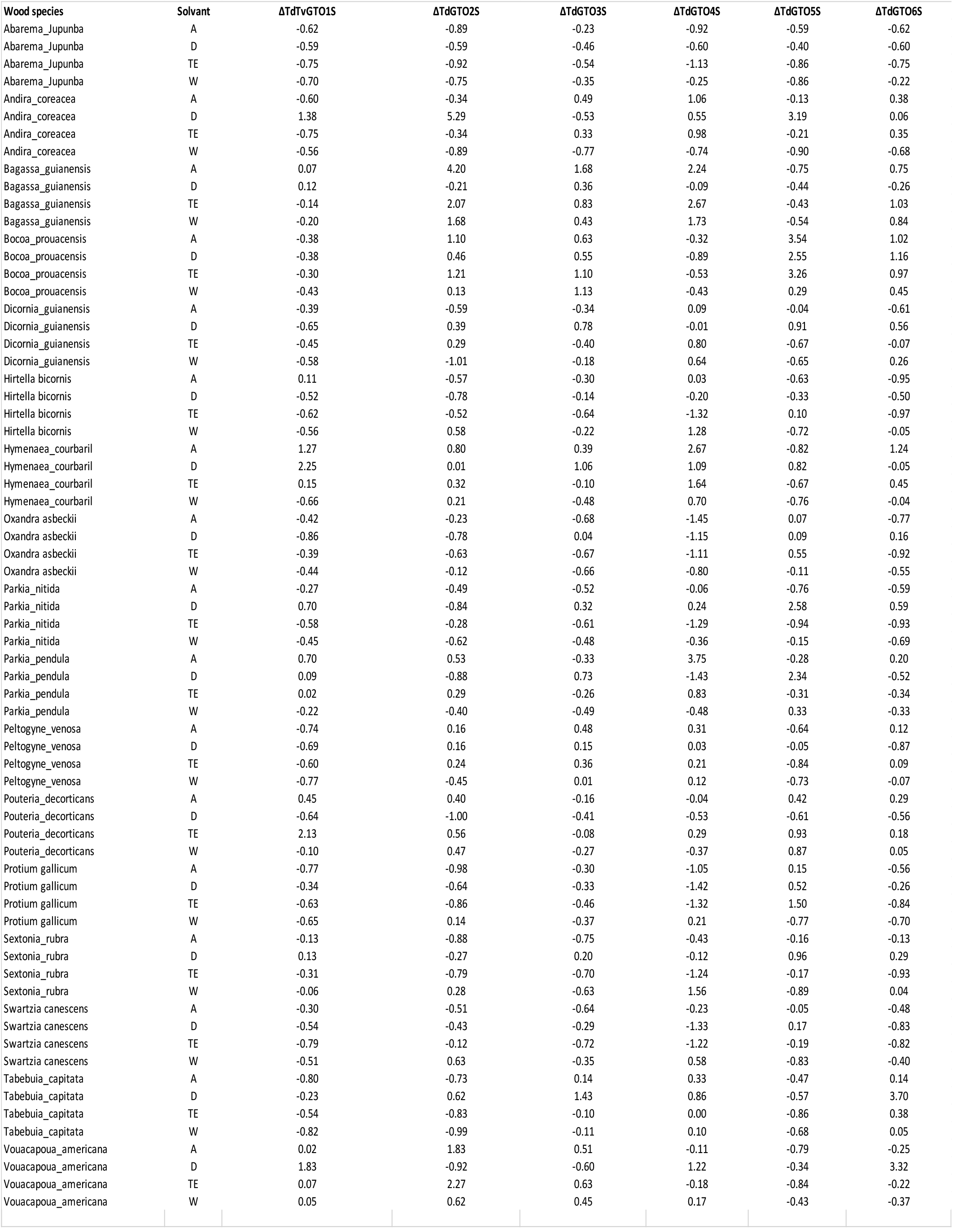
Assays was performed as described in (Deroy et al., 2015). The experimental procedure was performed in 96-well microplate (Harshell, Biorad) and the measurements were carried out with real time PCR detection system (CFX 96 touch, Biorad). The assays were achieved as follows: 5 μL of Tris-HCl (150 mM) pH 8.0 buffer, 2 μL of wood extracts at an initial concentration of 1 mg.mL^−1^ in DMSO, 2 μL of proteins (final concentration of either 10 or 20 μM depending on the corresponding assays), 2 μL of SYPRO^®^ orange diluted 62 fold (Sigma) and 14 μL of ultra-pure water. The microplate was centrifuged 30 s at 4000 *g*. The fluorescence was measured (excitation at 485 nm and emission at 530 nm) each minute starting with 3 min at 5 °C and increasing temperature from 5 to 95 °C with a step of 1 °C.min^−1^. The denaturation temperature (T_d_), which corresponds to the temperature where the protein is 50% unfolded, was determined using the first derivative of the obtained data in the presence or in the absence of potential ligands. As reference, experiments were conducted by adding DMSO only, allowing the determination of T_d_ ref. The corresponding values are the average of three technical repetitions, standard deviation remaining in all cases below 10%. Then, the difference between the denaturation temperature of the protein incubated with wood extracts and with DMSO only (T_d_ ref) were calculated in order to obtain the thermal shift (ΔT_d_). Absolute values of ΔT_d_ were then reduced centred. Solvent : A: Acetone; D: Dichloromethane; TE: Toluene/Ethanol; W: Water

